# Jumbo phages are active against extensively-drug-resistant eyedrop-associated *Pseudomonas aeruginosa* infections

**DOI:** 10.1101/2023.05.08.539869

**Authors:** Ana Georgina Cobián Güemes, Pooja Ghatbale, Alisha N. Blanc, Chase J. Morgan, Andrew Garcia, Jesse Leonard, Lina Huang, Grace Kovalick, Marissa Proost, Megan Chiu, Peiting Kuo, Joseph Oh, Smruthi Karthikeyan, Rob Knight, Joe Pogliano, Robert T. Schooley, David T. Pride

## Abstract

Antibiotic resistant bacteria present an emerging challenge to human health as the pressure instituted on the microbial world through the liberal use of antibiotics has resulted in their emergence across the globe. Those bacteria that acquire mobile genetic elements such as plasmids are especially concerning because those plasmids may be shared readily with other microbes that then can also become antibiotic resistant. Serious infections have recently been related to contamination of preservative-free eyedrops with extensively drug resistant (XDR) isolates of *Pseudomonas aeruginosa*, already resulting in three deaths. These drug-resistant isolates cannot be managed with most conventional antibiotics. We sought to identify alternatives to conventional antibiotics for lysis of these XDR isolates, and identified multiple bacteriophages (viruses that attack bacteria) that killed them efficiently. We found both jumbo phages (>200kb in genome size) and non-jumbo phages that were active against these isolates, the former killing more efficiently. Jumbo phages effectively killed the 3 separate XDR *P. aeruginosa* isolates both on solid and liquid medium. Given the ongoing nature of the XDR *P. aeruginosa* eyedrop outbreak, the identification of phages active against them provides physicians with several novel potential alternatives for treatment.

## INTRODUCTION

Antibiotic resistant bacteria present a serious and growing challenge as their numbers continue to increase across the globe (Murray et al. 2022). Most notable amongst those organisms whose propensity for antibiotic resistance has posed a challenge are the ESKAPE (*Enterococcus faecium*, *Staphylococcus aureus*, *Klebsiella sp.*, *Acinetobacter sp.*, *Pseudomonas aeruginosa*, and *Enterobacter sp.*) pathogens, which are responsible for the majority of nosocomial infections worldwide (Ma et al. 2020) These particular microbes have the potential to be MDR (multi-drug resistant; resistant to at least one antimicrobial in 3 different classes of antimicrobials) or XDR (extensively drug resistant; resistant to almost all approved antimicrobials), and sometimes require the use of lesser-used antibiotics for treatment or antibiotic alternatives when they cause infections in humans (Sundermann et al. 2023) A recent outbreak of XDR *P. aeruginosa* contaminating eye drops in the U.S. is a perfect example of the threat that such microbes may pose to the population long-term (Shoji et al. 2023; Morelli, Kloosterboer, and Omar 2023). Thus far, there have been at least 3 fatalities secondary to these *P. aeruginosa* isolates, which are incredibly difficult to treat with conventional antibiotics (CDC, https://www.cdc.gov/hai/outbreaks/crpa-artificial-tears.html).

Many bacteria are capable of acquiring new resistance genes through the process of conjugation where they may acquire mobile genetic elements such as plasmids (Partridge et al. 2018) In the current outbreak, *P. aeruginosa* has acquired the VIM beta lactamase, which confers resistance to carbapenem antibiotics amongst others (Meletis et al. 2014).This renders the organism very difficult to treat with antibiotics, and because of the concern that it may be mobile, also raises significant concern for spread of the carbapenemase to other bacteria.

Bacteriophages (viruses that attack bacteria) represent a rising therapeutic option for the treatment of antibiotic resistant bacteria (Strathdee et al. 2023). Mechanisms that account for reduced susceptibility of bacteria to antibiotics generally do not affect their susceptibilities to phages (Burmeister et al. 2020). Interest in the clinical use of phages for the treatment of drug resistant bacterial infections has been kindled by favorable outcomes reported in an increasing number of case reports and case series (Waters et al. 2017; Dedrick et al. 2019). In view of the ongoing eyedrop-associated *P. aeruginosa* outbreak in the U.S., we examined a collection of anti-*Pseudomonas* phages against 3 XDR *P. aeruginosa* isolates that were obtained from patients affected by this outbreak to identify phages capable of killing them and to identify whether they may be capable of inhibiting the XDR isolates across different conditions.

## RESULTS

### Pseudomonas aeruginosa isolates

We obtained each of the 3 separate *P. aeruginosa* isolates responsible for the ongoing outbreak found in eyedrops (Sundermann et al. 2023) from the CDC/FDA AR Isolate Bank (https://wwwn.cdc.gov/ARIsolateBank/; PA747, PA748, PA749, corresponding IDS in the AR bank are: 1268, 1269 and 1270). Antimicrobial susceptibility testing by microbroth dilution in the UC San Diego Center for Advanced Laboratory Medicine Clinical Microbiology Laboratory confirmed their XDR status (**Table 1**). The isolates were highly resistant to most beta-lactam antibiotics, including ceftolozane/tazobactam and ceftazidime/avibactam. Since the organisms possess a carbapenemase, treatment with meropenem and piperacillin/tazobactam would not be recommended despite intermediate MIC values in microbroth dilution assays. The only antibiotic to which the microbes demonstrated reproducible susceptibility was colistin (Sabnis et al. 2021).

**Table 1.** Antibiotic susceptibility of *Pseudomonas aeruginosa* epidemic isolates. MIC values are shown in ug/ml. Table is colored by interpretation, where dark gray represents resistant, mid gray represents intermediate MICs and clear represents susceptible MICs.

| Antibiotic | PS747 | PS748 | PS749 |
| --- | --- | --- | --- |
| Aztreonam | >16 | >16 | >16 |
| Ceftazidime | >16 | >16 | >16 |
| Cefepime | >16 | >16 | >16 |
| Meropenem | >8 | 8<sup>a</sup> | 8<sup>a</sup> |
| Piperacillin/Tazobactam | 64 | >64 | 32<sup>a</sup> |
| Ceftolozane/Tazobactam | >8/4 | >8/4 | >8/4 |
| Ceftazidime/Avibactam | >16/4 | >16/4 | >16/4 |
| Ciprofloxacin | >2 | >2 | >2 |
| Levofloxacin | >4 | >4 | >4 |
| Gentamicin | >8 | >8 | >8 |
| Tobramycin | >8 | >8 | >8 |
| Amikacin | >32 | >32 | >32 |
| Colistin | ≤2 | ≤2 | ≤2 |

<sup>a</sup>Represents intermediate MIC values

We also sequenced each of the *P. aeruginosa* isolates obtained from the CDC in this study. Our results were largely consistent with sequencing of these same reported by other groups (Sundermann et al. 2023). We assembled the genome sequences through a combination of Illumina short-read and Nanopore long-read sequencing. We confirmed that each of the genomes was generally roughly 7Mbp in size, with nearly 66% G+C content (**Table S1**). Although we were able to sequence the genomes to near completion, we were not able to assemble the genomes below 25 separate 100 Kbp contigs. Many putative antibiotic resistance genes were identified in the genome sequences, including those associated with resistance to beta lactam, chloramphenicol, fosfomycin, macrolides, aminoglycosides, and fluoroquinolone antibiotics (**Figure 1, Panel C**). The VIM beta-lactamase sequence was identified in each of the organisms. Notably a contig of 5 Kbp carries 2 beta-lactamases (VIM-2 and OXA-10), two aminoglycoside resistance genes and a transposase (**Figure 1, Panel A**). A plasmid of 78 Kbp carries a transposase, two beta-lactamases (GES and CatB) and several other antibiotic resistance genes (**Figure 1, Panel B**).

**Figure 1.**
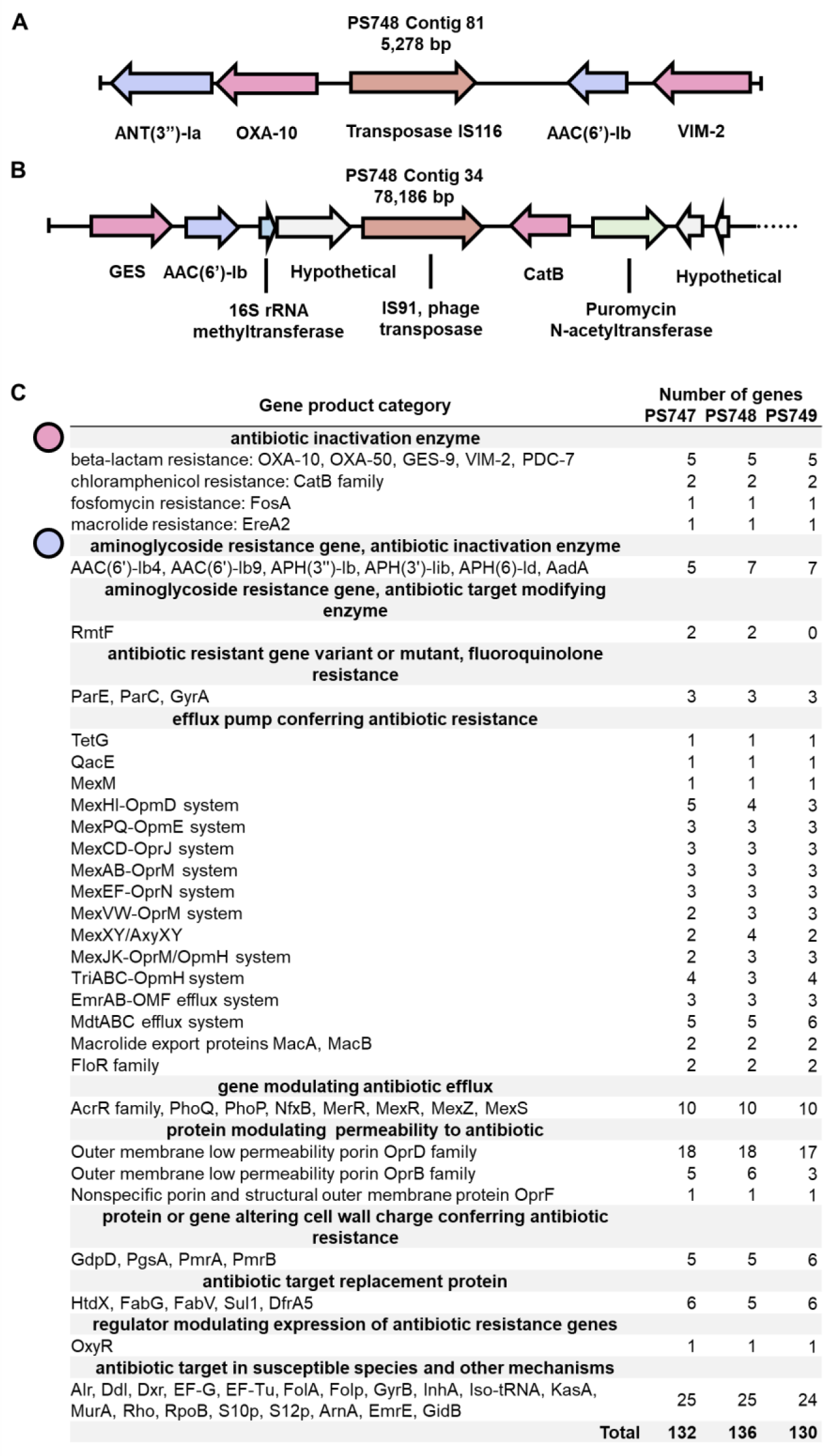
Antibiotic resistance genes in eyedrop-associates *Pseudomonas aeruginosa* genomes. A) Selected IS element of PS748. B) Selected plasmid fragment in PS748. C) Antibiotic resistance genes in PS748. Genome annotations were performed in the BV-VCR server using the CARD database.

### Bacteriophages active against XDR isolates

We tested a collection of phages to determine whether some were active against the XDR *P. aeruginosa* isolates and other recently identified clinical *P. aeruginosa* isolates (**Table S2**). We found 14 separate phages with activity against one or more of the *P. aeruginosa* isolates (**Figure 2**). We first characterized the phage-host interactions using standard plaque assays on solid media. Two of the *P. aeruginosa* isolates were not lysed by any of the 14 phages in the collection. One or more phages exhibited lytic activity against each of the other 12 isolates. Ten of the 14 phages exhibited lytic activity against one or more of the eyedrop-associated isolates but only three phages produced clear plaques at the highest dilution tested (**Figure 2**). The jumbo phages (tailed phage genomes >200kb) had significant lytic activity against the eyedrop isolates. These jumbo phages included the previously described PhiKZ (Krylov et al. 2021) and PhiPA3 (Monson et al. 2011), but also 2 additional phages, ANB1 and PhiPizzaParty that were found in this study.

**Figure 2.**
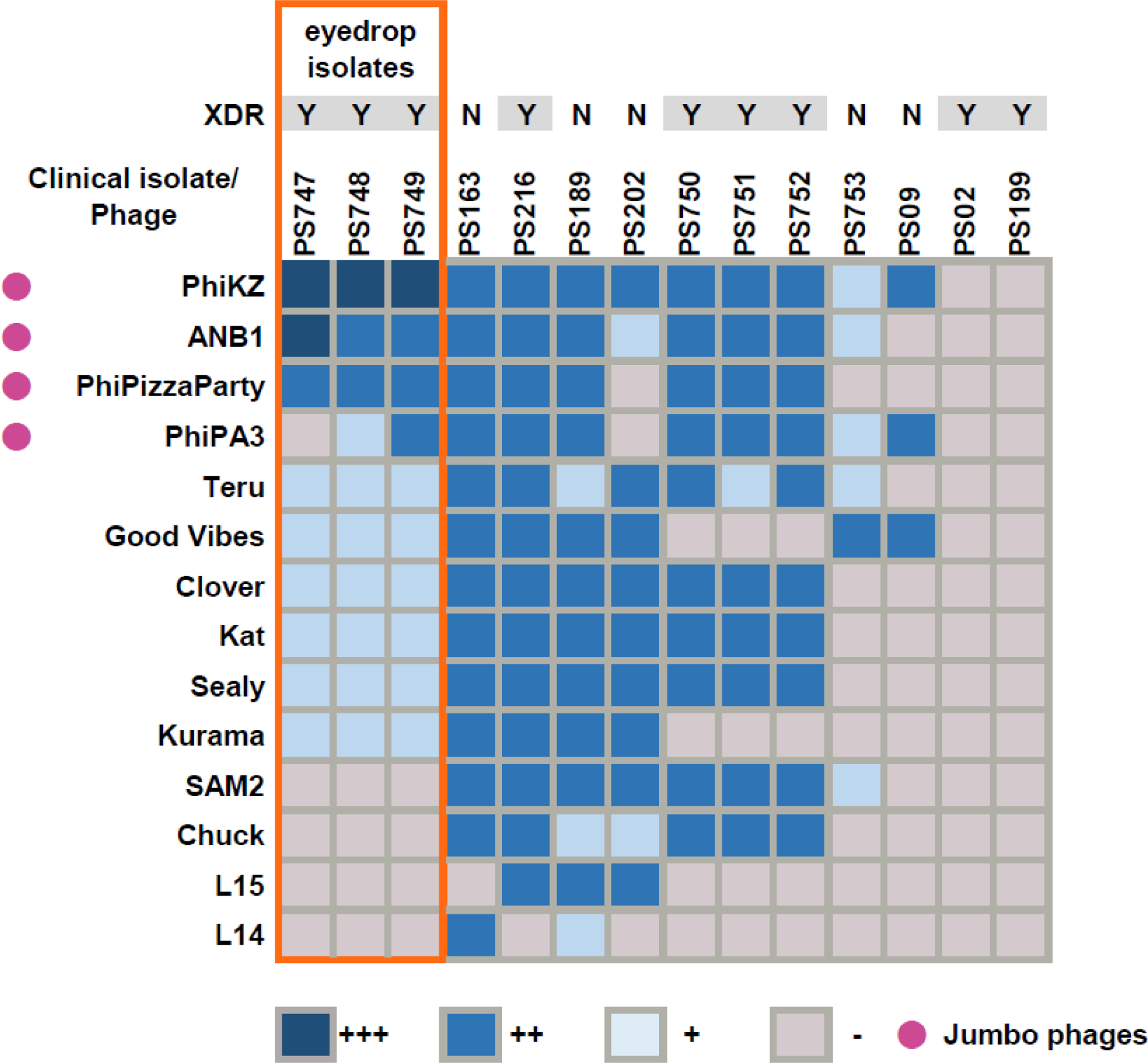
Host range of phages in selected *P. aeruginosa* clinical isolates. +++ lysis at 1×10^−6^, ++ lysis at 1×10^−5^ to 1×10^−3^, + lysis at 1×10^−2^, - lysis at undiluted

### Genome characterizations and imaging of phages

We characterized the genome and structures of the jumbo phages in this collection that exhibited lytic activity against the outbreak-associated XDR *P. aeruginosa* isolates. Phages PhiPizzaParty and ANB1 were similar in size and gene content to PhiKZ (**Figure 3**). They had many similarities and some differences along their genome structures, indicating that they were likely derived from similar ancestral phages (**Figure 3**). We did not identify any gene content to suggest they may be involved in lysogenic infections. The closest relative to PhiPizzaParty and ANB1 was phage SL2 (Latz et al. 2017), these phages have an identity greater than 97% between each other. They are substantially different from phage PA7 (NC_042060.1) and PhiPA3 (Monson et al. 2011) (**Supplemental Figure 1**). We found that phage PhiPizzaParty was 98.4% similar to PhiKZ, with most of the differences arising in the gene structures of endonucleases, structural head protein, tail fibers and other proteins without annotation. Phage ANB1 also was 98% similar to PhiKZ, with differences arising in similar gene structures. ANB1 was more divergent from PhiKZ than PhiPizzaParty (**Figure S1**). The genomes for non-jumbo phages were also sequenced (**Figure S2**).

**Figure 3.**
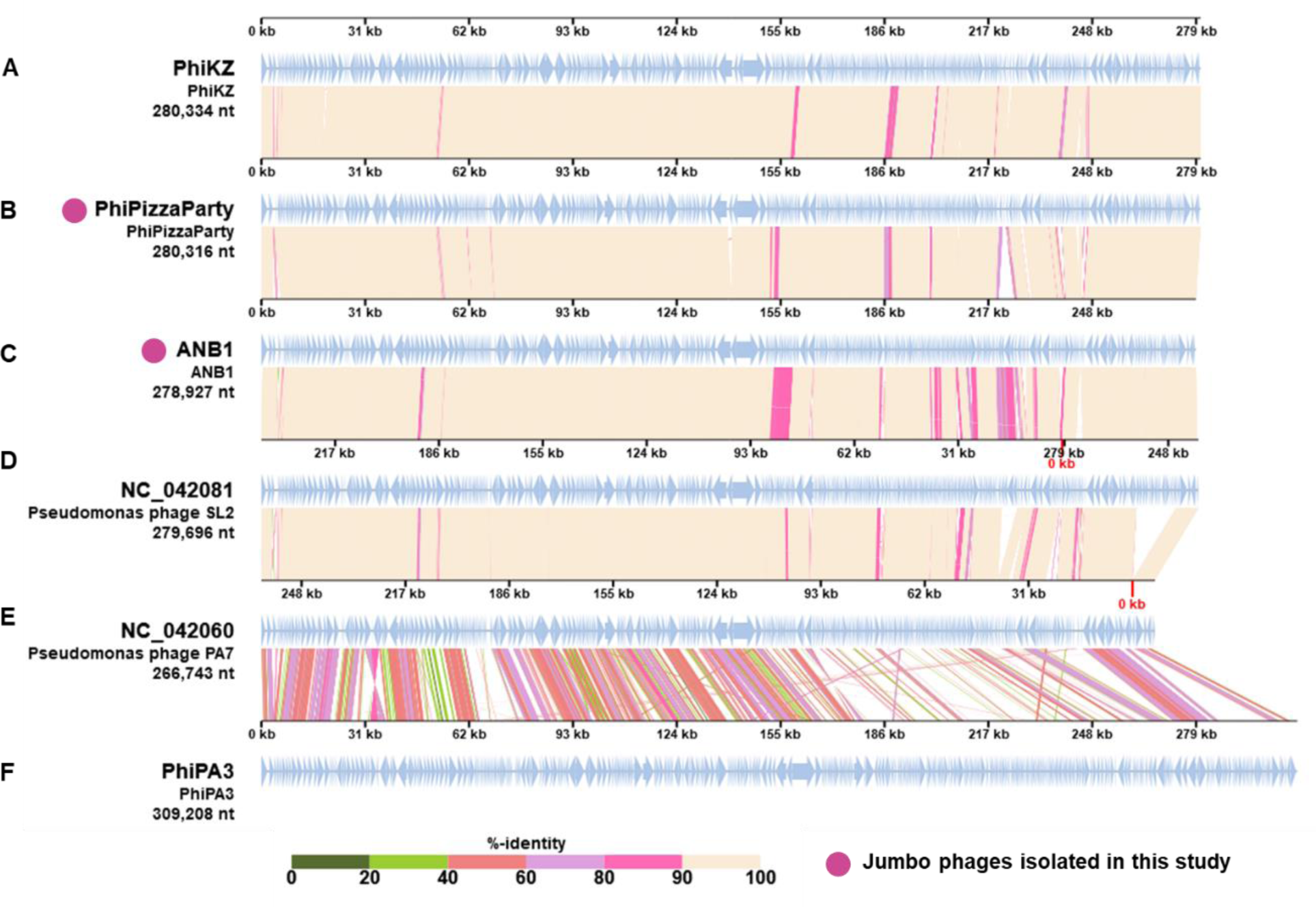
Comparative genomics of Jumbo phages. Jumbo phages tested against XDR *Pseudomonas aeruginosa* eyedrop isolates (A, B, C, F) and its closest relatives (D, E). Jumbo phages PhiPizzaParty (B) and ANB1 (C) were discovered in this study. Comparative genomics were performed in the VIP tree server.

We previously demonstrated that many *Pseudomonas* jumbo phages including PhiKZ replicate by enclosing their genome within a proteinaceous shell, forming a structure (phage nucleus) that segregates phage DNA from the host cell cytoplasm (Chaikeeratisak, Nguyen, Khanna, et al. 2017; Chaikeeratisak, Nguyen, Egan, et al. 2017; Knipe et al. 2022) Nucleus forming phage are particularly well suited for phage therapy because they are broadly immune to many bacterial phage defense systems (Mendoza et al. 2020; Malone et al. 2019) The gene that encodes the nuclear shell protein, Chimallin (Prichard et al. 2023; Laughlin et al. 2022) in PhiKZ and other nucleus forming phages was present in both PhiPizzaParty and ANB1. To determine whether these phages also formed a phage nucleus, we performed fluorescence microscopy on phage infected cells (**Figure 4, Panels O-P**). Imaging of DAPI-stained infected cells showed that, similar to PhiKZ, PhiPizzaParty degrades the host chromosome and centers its genome in the cell in a manner that is consistent with nucleus-forming phage. Together, this supports the conclusion that PhiPizzaParty and probably ANB1 are nucleus-forming phages.

**Figure 4.**
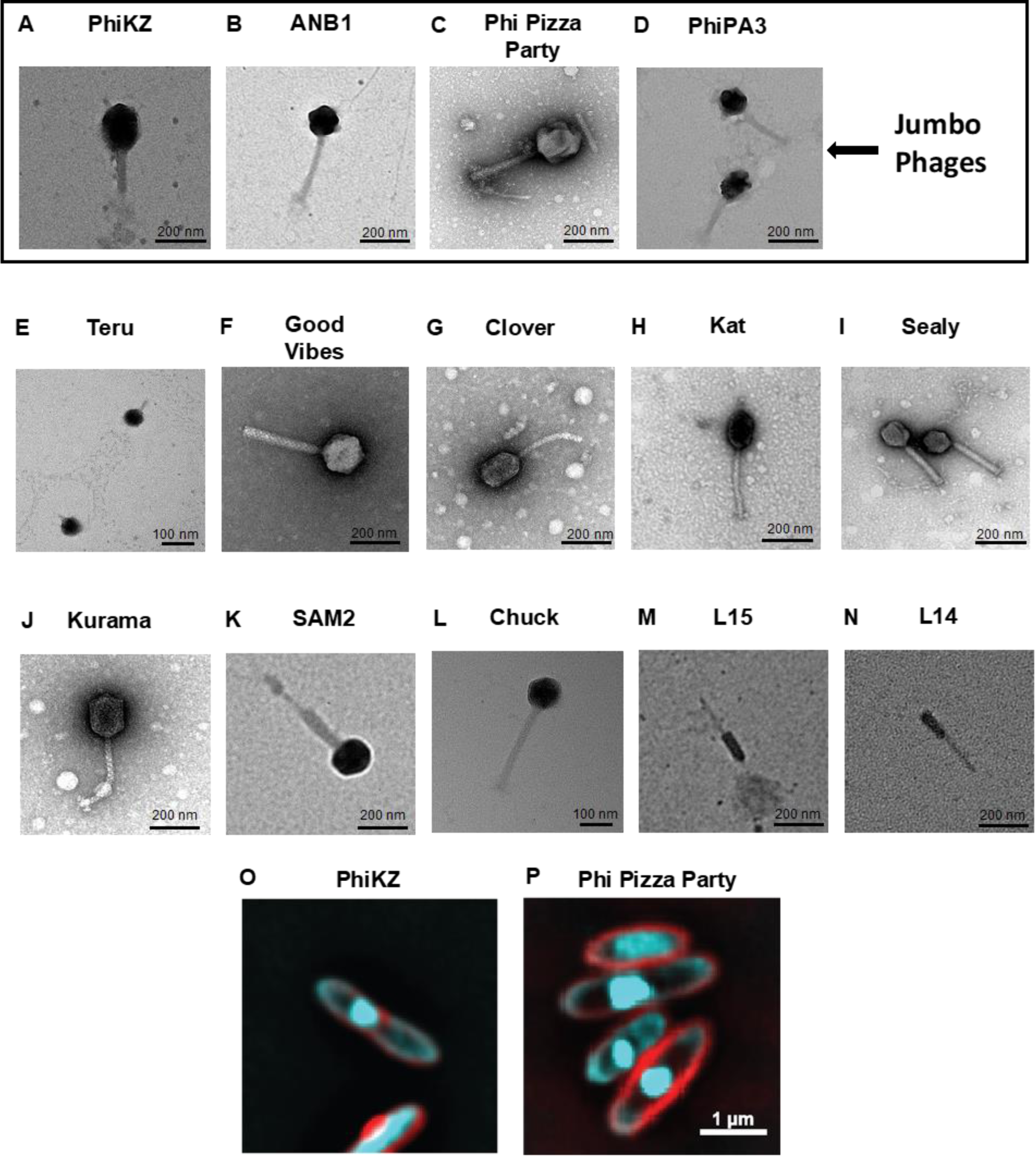
TEM of phages used in this study. A to D) jumbo phages. E to N) non jumbo phages. O and P) fluorescence microscopic images of *Pseudomonas* aeruginosa infected with phage PhiKZ and Phi Pizza Party respectively. The cell membrane is stained with FM4-64 (Red) and the DNA is stained with DAPI (Cyan). Phage nucleus can be seen centered in the cell.

### TEM analysis

We imaged each of the phages in this study using Transmission Electron Microscopy to determine further their structures. The images of each of the phages demonstrate that most have an icosahedral head structure with relatively long, noncontractile tail structures as are often observed in the family *Siphoviridae* (**Figure 4, Panels A-N**) (Ackermann, HW. 2009). Phage Teru, which also was active against many of the *P. aeruginosa* isolates was the most notable exception with more of a tail stub similar to that observed in the family *Podoviridae* (**Figure 4, Panel E**).

### Efficiency of plating analysis

To further characterize the activity of the phages against the XDR *P. aeruginosa* isolates, we performed an Efficiency of Plating (EOP) analysis. We tested 7 of the most active phages from the plaque assay analysis (**Figure 2**) and found that each of them was also active in an EOP analysis (**Table 2**). Most of the non-jumbo phages produced relatively high EOP values, with phages Good Vibes, Sealy, and Clover producing EOP values well above 10^−3^ (**Table 2**). Phage Teru was the only non-jumbo phage to produce EOP values similar to those of the jumbo phages. The jumbo phages produced EOP values above 10^−3^ in most cases, with PhiKZ and ANB1 producing values ≥10^−3^ for each of the XDR isolates. Phage PhiPizzaParty produced EOP values ≥10^−3^ for isolates PS748 and PS749, but not for PS747.

**Table 2.** Efficiency of plating.

| Phage | Titer on PA01 | Titer on PS747 | Titer on PS748 | Titer on PS749 | EOP PS747 | EOP PS748 | EOP P749 |
| --- | --- | --- | --- | --- | --- | --- | --- |
| PhiKZ | $1 \times 10^{10}$ | $1.6 \times 10^8$ | $2 \times 10^8$ | $2.4 \times 10^8$ | **$1.6 \times 10^{-2}$** | **$2 \times 10^{-2}$** | **$2 \times 10^{-2}$** |
| ANB1 | $6 \times 10^9$ | $1.3 \times 10^8$ | $4 \times 10^7$ | $2 \times 10^7$ | **$2 \times 10^{-2}$** | **$1 \times 10^{-1}$** | **$3 \times 10^{-3}$** |
| PhiPizzaParty | $7.3 \times 10^9$ | $1.4 \times 10^5$ | $4 \times 10^7$ | $1 \times 10^8$ | $2 \times 10^{-5}$ | **$1 \times 10^{-1}$** | **$1 \times 10^{-1}$** |
| Teru | $2 \times 10^{10}$ | $2 \times 10^4$ | $2 \times 10^4$ | $2 \times 10^4$ | $1 \times 10^{-6}$ | **$1 \times 10^{-1}$** | **$1 \times 10^{-1}$** |
| Clover | $2 \times 10^{11}$ | $2 \times 10^3$ | $2 \times 10^3$ | $1 \times 10^0$ | $1 \times 10^{-8}$ | $1 \times 10^{-8}$ | $5 \times 10^{-12}$ |
| Sealy | $2.4 \times 10^{10}$ | $8 \times 10^5$ | $1 \times 10^0$ | $1 \times 10^0$ | $4 \times 10^{-5}$ | $5 \times 10^{-11}$ | $5 \times 10^{-11}$ |
| Good Vibes | $1 \times 10^{11}$ | $1 \times 10^0$ | $1 \times 10^0$ | $1 \times 10^0$ | $1 \times 10^{-11}$ | $1 \times 10^{-11}$ | $1 \times 10^{-11}$ |

EOP values above $1 \times 10^{-3}$ are considered effective.

### Liquid media suppression

We sought to determine whether our jumbo phages also might be active in liquid medium in addition to the solid medium against the XDR *P. aeruginosa* isolates. There was substantial inhibition of each of the XDR isolates at up to 24 hours of co-cultivation (**Figure 5**). For example, there was significant inhibition of isolate PS747 when cultivated with most of the phages, particularly when phage cocktails were used (**Panel A**). Isolate PS748 was inhibited by the phages in liquid medium but was more variable than was observed for PS747 (**Panel B**), and PS749 had its growth inhibited substantially by some of the phages (**Panel C**).

**Figure 5.**
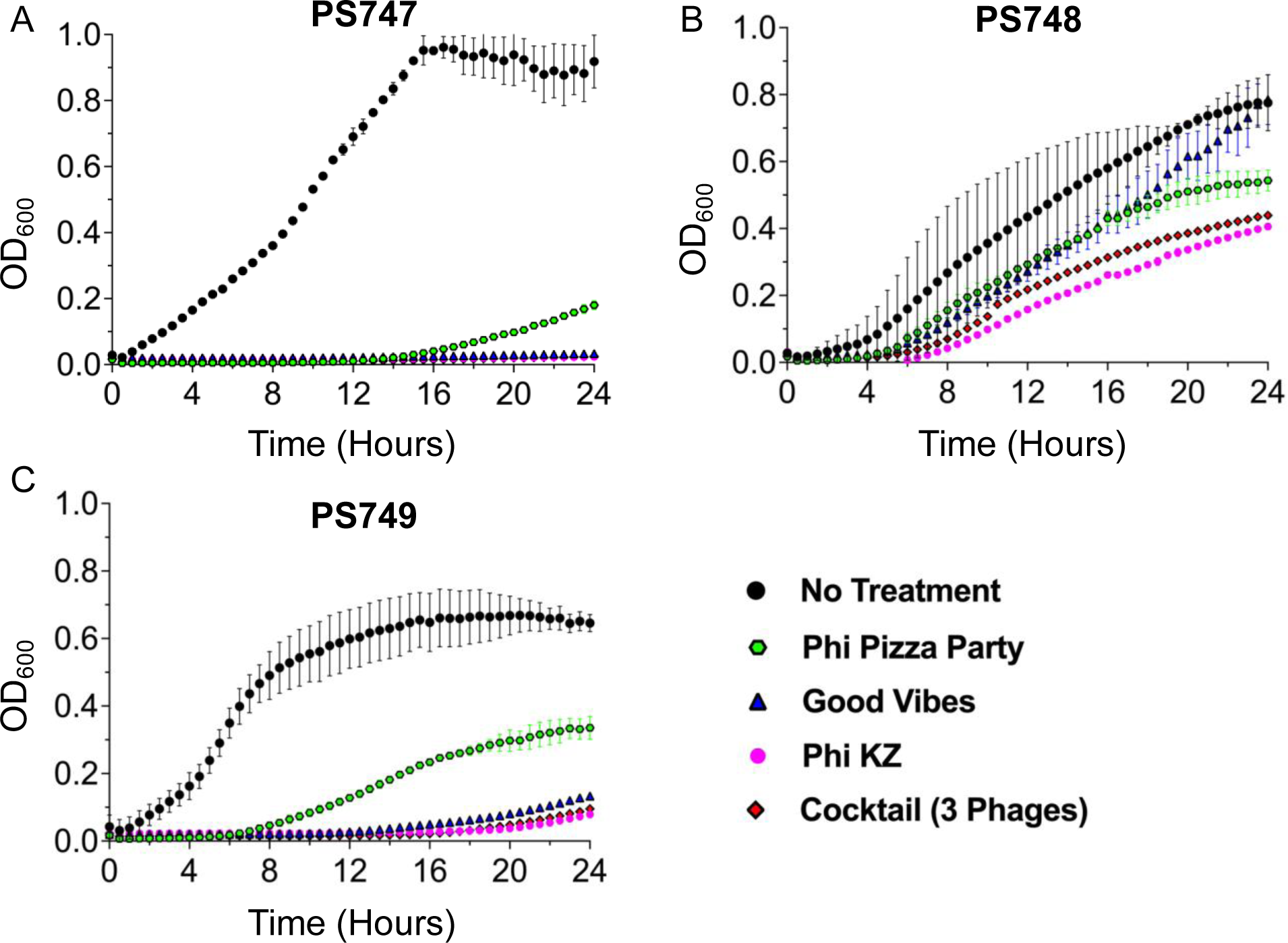
Inhibition of bacteria growth by phages in broth medium.

## DISCUSSION

We identified three jumbo phages with significant activity against three XDR *P. aeruginosa* isolates associated with a recently described multi-state outbreak among users of ophthalmic eye drops on solid and in liquid medium. We also identified a non-jumbo phage with significant activity on solid medium. While we do not have specific information on some of these phages to describe their receptors and interactions with their host *P. aeruginosa* strains, the jumbo phages are similar to previously used phages such as OMKO1 (Chan et al. 2016), which already have been used rather extensively in phage therapy applications. The phages we identified in this study (ANB1, PhiPizzaParty, and Teru) were all identified from wastewater on the UCSD campus, strongly suggesting that these phages are already circulating through the population.

The use of jumbo phages that have nucleus-like structures may have certain advantages compared to more conventional bacteriophage therapies, as the nuclear shell excludes many DNA-targeting anti-phage systems (Mendoza et al. 2020; Malone et al. 2019; Nguyen et al. 2021). PhiKZ-like phages also degrade the host chromosome early in infection further decreasing the likelihood of lysogenic infection (Kraemer et al. 2012; Erb et al. 2014).

It is important to note that carbapenems are often the last line of therapy for the treatment of many pathogens but is not necessarily the case for *P. aeruginosa*. This is because the organism can have multiple different mechanisms by which it can resist carbapenems (Pai et al. 2001), which may persist after the organism were to be cured of the VIM beta lactamase. In many cases of *P. aeruginosa*, aminoglycoside antibiotics such as amikacin represent the last line of therapy, but in this case, each of the isolates were already resistant to aminoglycosides. These isolates present medical conundrums for therapy, as they are resistant to most commonly used antibiotics, and by the time most laboratories would be able to test for susceptibility to secondary antibiotics such as colistin and cefidericol, the infection could become much more difficult to eradicate. The knowledge that PhiKZ (a widely available phage) has significant activity against these isolates could make it a candidate for use in phage therapy applications against these XDR *P. aeruginosa* isolates.

One of the primary concerns with the use of phages to treat bacterial infections is the potential that phages may integrate into the host genomes and cause long-term phage infections rather than immediate lysis. We did not identify any genes in either PhiPizzaParty, ANB1, or Teru that would suggest they have the potential for lysogenic infections. PhiKZ-like phages also degrade the host chromosome early in infection further decreasing the likelihood of lysogenic infection (Kraemer et al. 2012) Given the immense concern for the continued spread of these XDR isolates with the use of contaminated eye drops, and the relatively few antibiotics that may be effective, development/purification of phages such as those identified here could offer critical treatment alternatives to eliminate the ongoing risk from these XDR *P. aeruginosa* strains.

## METHODS

### Bacterial strains, Bacteriophages, and culture conditions

*P. aeruginosa* strains used in the study (**Table S2**) were previously isolated from patients with pseudomonas infection at UCSD Centre for Advanced Laboratory Medicine. The antibiotic sensitivity testing was performed on the isolates using MicroScan NM46 and DNM2 panels. The phages used in this study were previously isolated from various environmental sources (**Table S3**). Luria Bertani (LB) media was used to grow the *P. aeruginosa* strains and phages at 37°C. While liquid phage/bacteria co-cultivation assays were performed in MacConkey media.

### Jumbo phage isolation

Wastewater was collected from the UCSD campus through the wastewater monitoring program (Karthikeyan et al. 2022; Fielding-Miller et al. 2023), and from several locations in southern California (**Table S3**). Jumbo phage isolation and purification was carried out as previously described (Saad et al. 2019). Wastewater was centrifuged at 3000xg for 10 min. The supernatant was collected and centrifuged at 4696xg for 55 min. The supernatant was removed, and the pellet was resuspended in 10 ml of SM buffer, this process was performed twice. The sample was treated with equal volume of chloroform to remove residual bacteria. Plaques were purified 3 times in 0.3% agar plates. Phages were produced by plate lysis in 0.1% agar.

### Host range evaluation and Efficiency of Plating (EOP)

*P. aeruginosa* isolates listed in (**Table S2**) were incubated overnight at 37°C. Next day, 100 µl of bacterial culture with an OD_600_ of 0.2 was mixed with 3 ml of melted LB soft agar (when soft agar temperature reaches ∼45°C). This mixture was then overlaid on LB Agar plate (1.5% Agar). A 5 μL sample of serially diluted phages and a 5 μL LB control were spotted onto the bacterial overlay, left for 30 min to dry, and then the plates were incubated overnight at 37°C. Next day, the number of plaques were counted. The EOP assays were carried out as previously described (Elizabeth Kutter 2009).

### Liquid phage/bacteria cultivation assays

The phage and bacterial co-cultivation assays were performed at MOI of 1. The bacterial cells from exponential phase were diluted to an OD_600_ of 0.1 in fresh LB broth. Each well within 96 well plate was inoculated with 20 µl of phage + 60 µl of bacteria (OD_600_ 0.1) and remaining volume of MacConkey media was added to a total volume of 200 µl. To evaluate three phage cocktails, one third volume of 20 µl of each phage was added. The OD_600_ was measured every 30 minutes at 37°C for 24 hours.

### Phage characterization using Transmission Electron Microscopy (TEM)

Carbon coated grid (PELCO SynapTek™ Grids, product# 01754-F) was placed on a drop of 10 µl of freshly made phage stock (∼10^10^ PFU/ml) and the grids were negatively stained with 2% uranyl acetate (pH 4.0) for 45 sec. Imaging was performed using Joel 1400 plus at University of California, San Diego - Cellular and Molecular Medicine Electron Microscopy Core (RRID:SCR_022039).

### Fluorescence Microscopy of Phage Infected *Pseudomonas aeruginosa*

PA01 K2733 was grown in LB to an OD of 0.5 and incubated with high titer phage lysate at a ratio of 1:100 lysate-to-culture at 30℃ with agitation for 30 min. FM4-64 dye was added to the cells at a final concentration of 6.5 ug/mL and then culture was spotted onto a 1% agarose 25% LB pads containing 0.6ug/mL DAPI and visualized on an Applied Precision DV Elite optical sectioning microscope with a Photometrics CoolSNAP-HQ2 camera (Applied Precision/GE Healthcare). Microscopic images were deconvolved using SoftWoRx v5.5.1. Image analysis and processing were performed in Fiji.

### Whole Genome Sequencing (WGS)

Total genomic DNA from the bacteriophages and bacterial isolates was extracted using QIAamp UltraSens Virus kit (Qiagen catalog number 53706) and DNeasy Blood & Tissue Kit (Qiagen catalog number 69504) respectively. Genome sequencing of bacteriophages and bacterial isolates was carried out using a paired end approach (2 × 150 bp) on iSeq100 and Miseq platform (Illumina) respectively. The sequences were assembled using de-novo assembly algorithm of CLC Genomics Workbench (CLC Genomics, Qiagen, version 9.5.3). Rapid Annotation using Subsystem Technology (RAST) pipeline (v2.0) was used to annotate the sequences (McNair et. al, 2018). For nanopore sequencing, bacterial DNA was extracted using the ZYMO DNA microprep kit. Libraries were constructed using the rapid sequencing kit and sequenced on the MinION platform.

## Abbreviations

PSA: *Pseudomonas aeruginosa*
XDR: extensively drug resistant
MDR: multi-drug resistant

## Declarations

### Availability of Data and Material

All sequences included in this study have been deposited in the NCBI Sequence Read Archive under BioProject accession.

### Author Contributions

Conceived and designed project: DTP and RTS.

Performed experiments: AGCG, PG, ANB, AG, JL, JO, MC, PK, and MP.

Analyzed the data: AGCG, PG, ANB, AG, DTP, and RTS.

Wrote and edited the manuscript: AGCG, PG, JP, DTP, and RTS.

Provided materials for the study: SK, RK.

All authors have reviewed the manuscript.

## Acknowledgements

We thank the UCSD Clinical Microbiology Laboratory for their participation in this work. We also thank the CDC for their generous sharing of the PSA isolates. We thank Dr. Elizabeth Villa and the EPI consortium for fruitful discussions.

## SUPPLEMENTAL DATA

**Figure S1.**
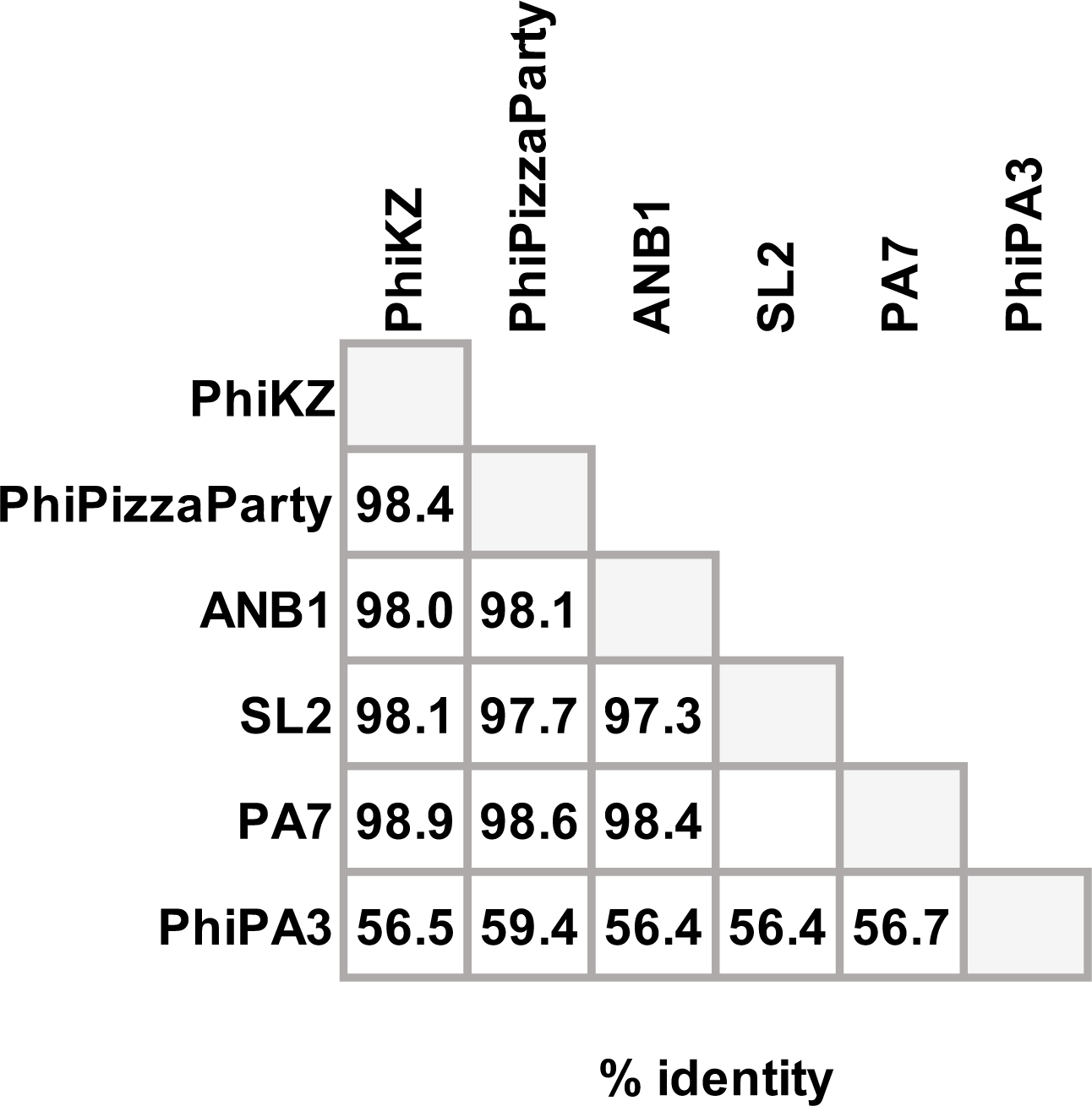
Identity amongst the jumbo phages presented in this work.

**Figure S2.**
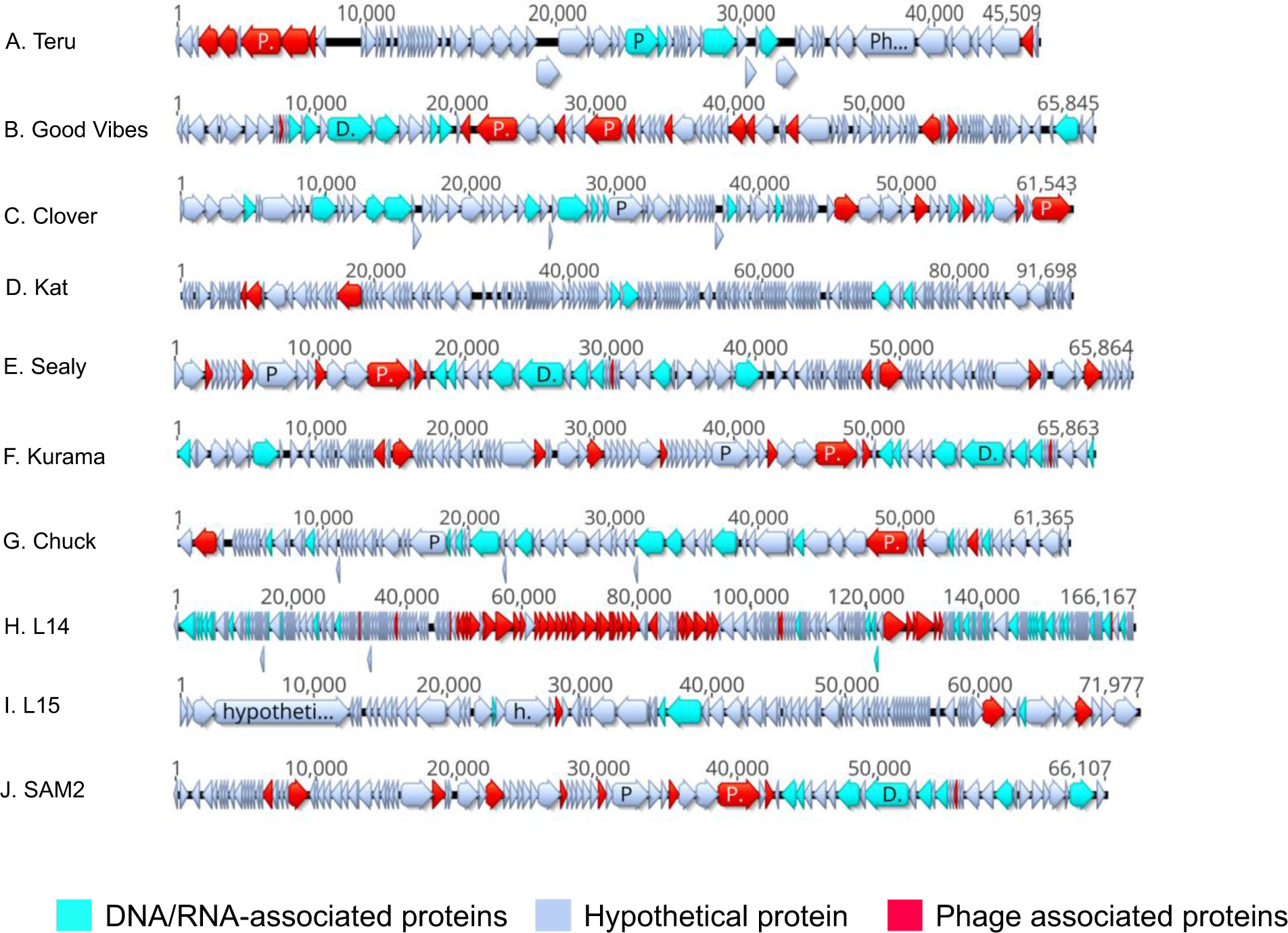
Genome annotation of non-jumbo phages characterized in the study.

**Table S1.**
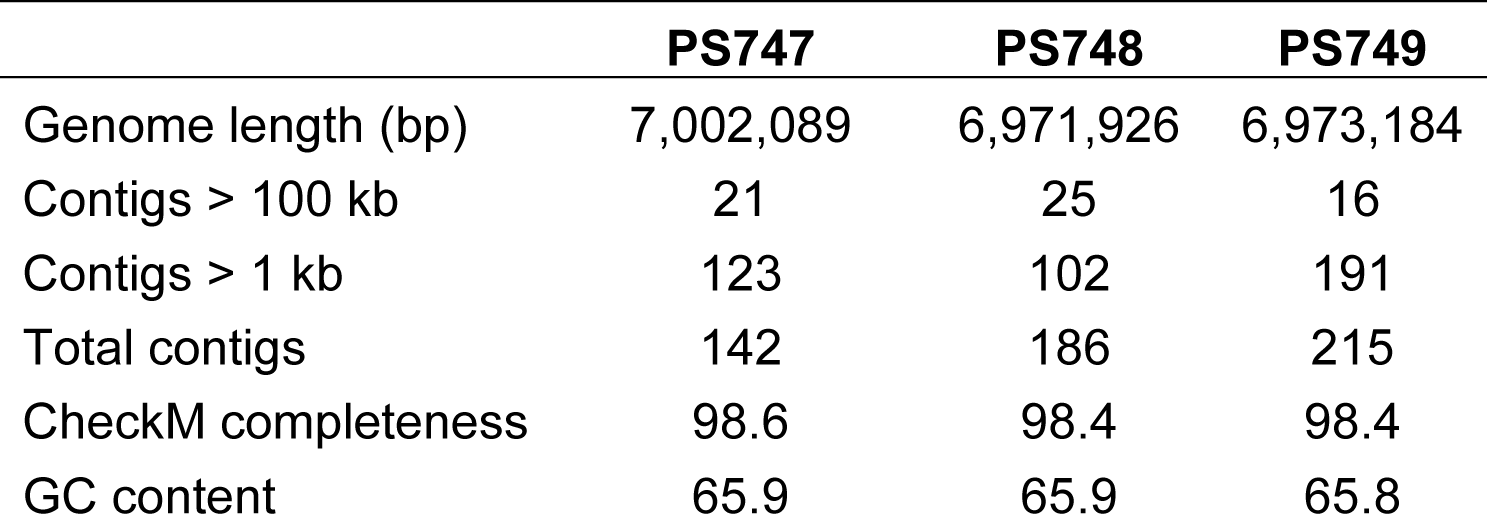
*P. aeruginosa* genomes sequenced in this study.

|  | **PS747** | **PS748** | **PS749** |
| --- | --- | --- | --- |
| Genome length (bp) | 7,002,089 | 6,971,926 | 6,973,184 |
| Contigs > 100 kb | 21 | 25 | 16 |
| Contigs > 1 kb | 123 | 102 | 191 |
| Total contigs | 142 | 186 | 215 |
| CheckM completeness | 98.6 | 98.4 | 98.4 |
| GC content | 65.9 | 65.9 | 65.8 |

**Table S2.**
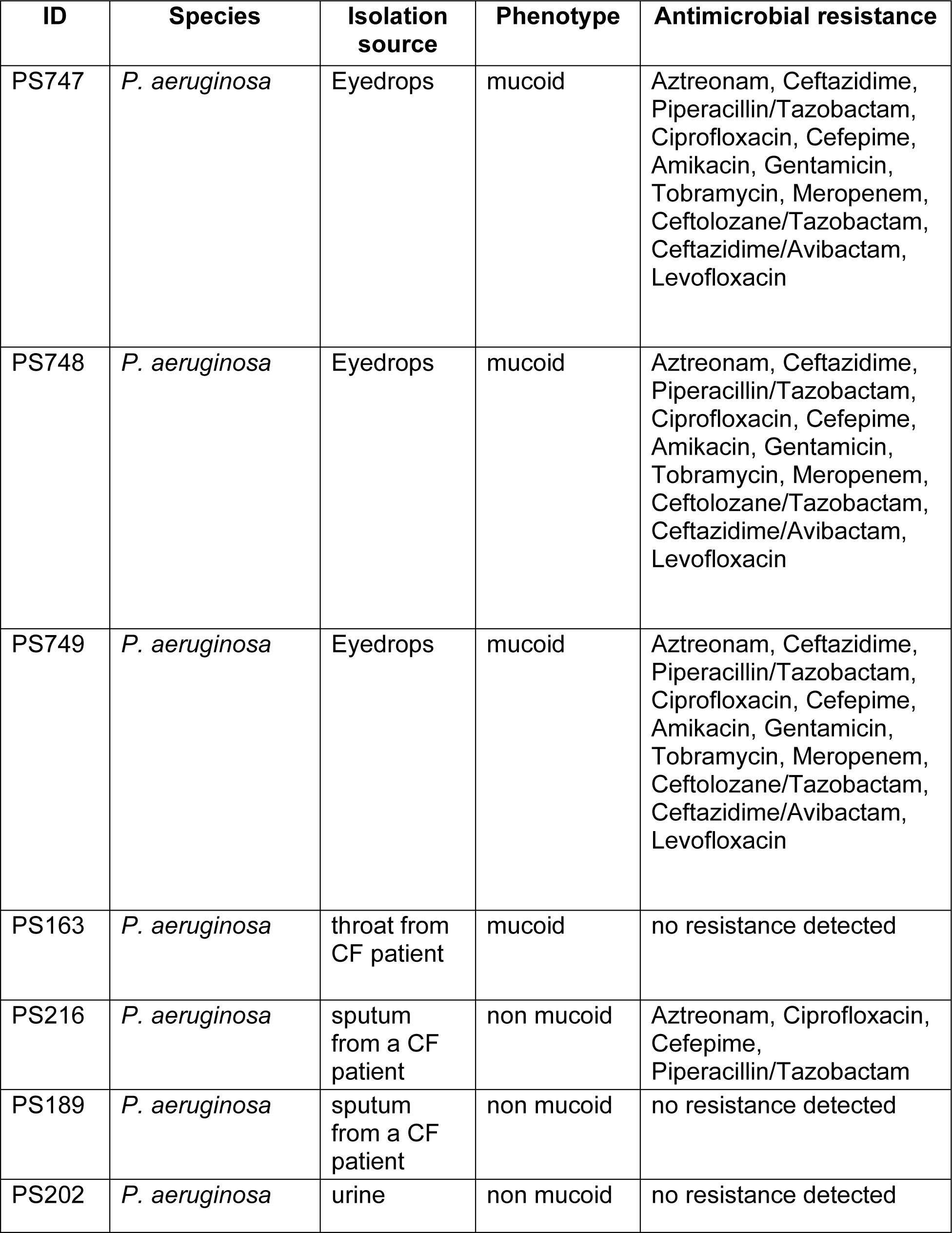
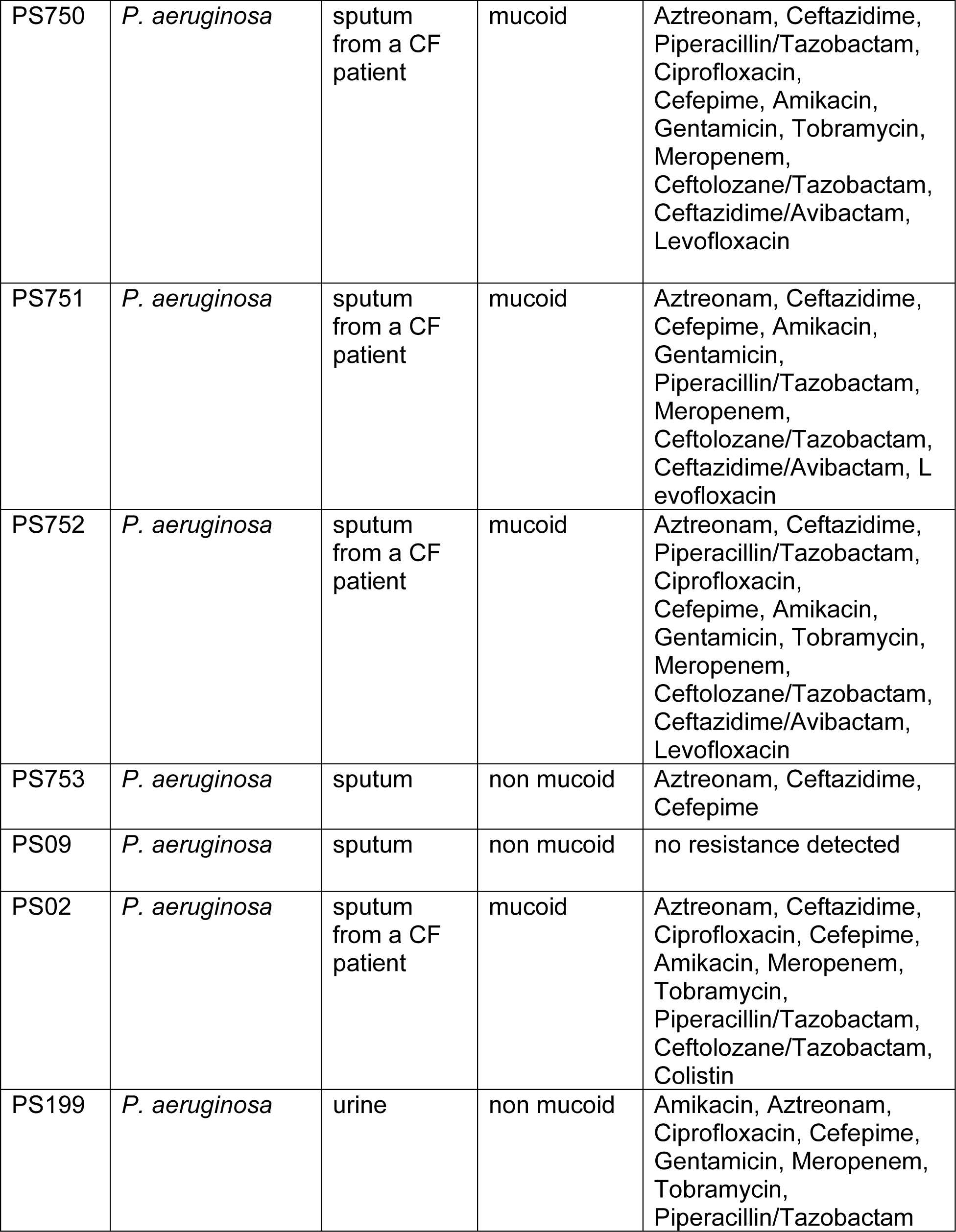
*P. aeruginosa* strains used in the study.

**Table S3.**
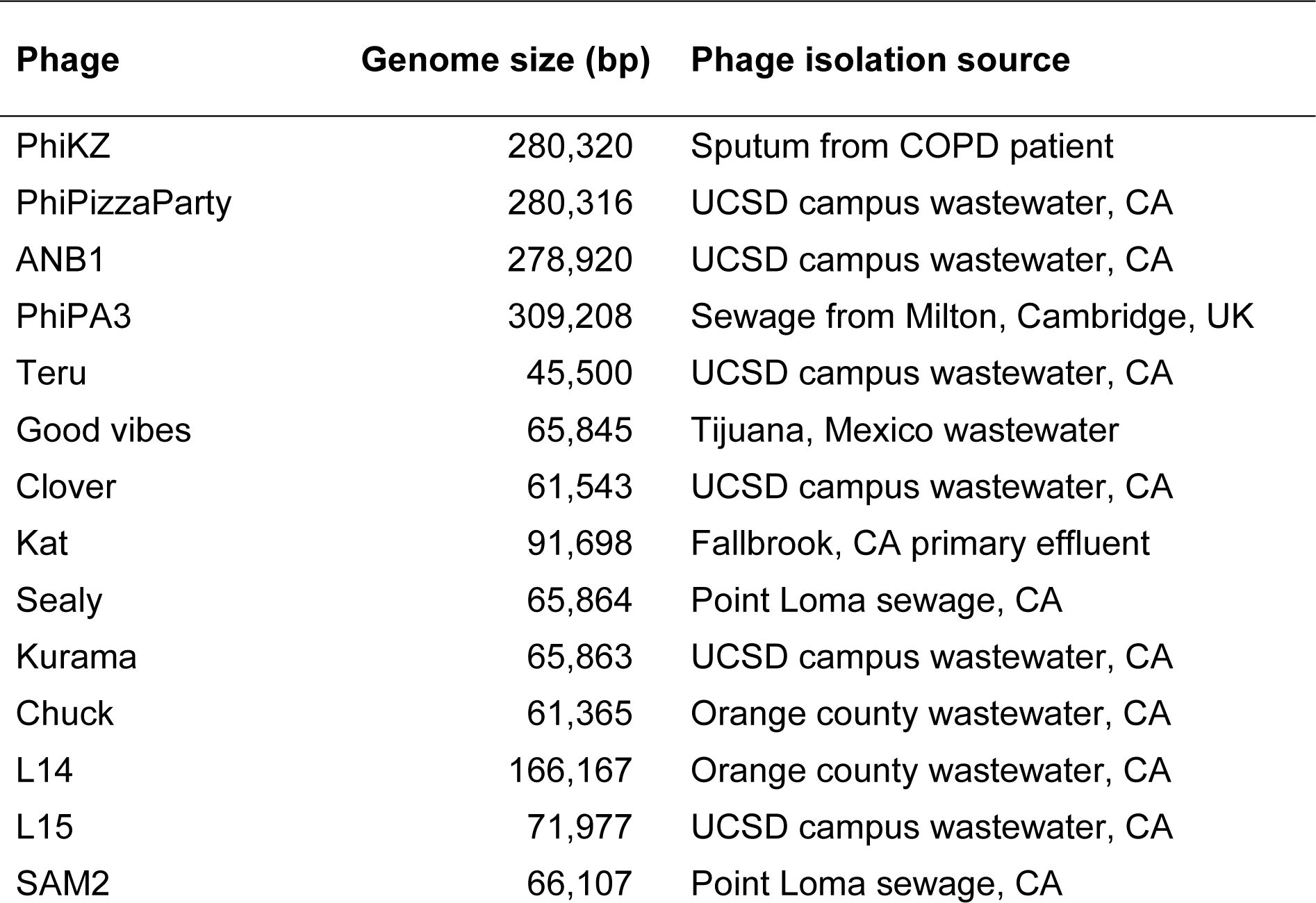
Bacteriophages lysis of *P. aeruginosa* epidemic isolates.

